# Neurons in the dorso-central division of zebrafish pallium respond to change in visual numerosity

**DOI:** 10.1101/2020.11.11.377804

**Authors:** Andrea Messina, Davide Potrich, Ilaria Schiona, Valeria Anna Sovrano, Scott E. Fraser, Caroline H. Brennan, Giorgio Vallortigara

## Abstract

Non-symbolic number cognition based on an approximate sense of magnitude has been documented in zebrafish. Here we investigated for the first time its neural bases. Zebrafish were habituated to a set of three or nine small dots associated with food reward. During habituation trials, the dots changed in their individual size, position and density maintaining their numerousness and overall surface area. In the dishabituation test, zebrafish faced a change (i) in number (from three to nine or vice versa with the same overall surface), (ii) in shape (with the same overall surface and number), or (iii) in size (with the same shape and number); in a control group (iv) zebrafish faced the same familiar stimuli as during the habituation. Using qPCR to measure modulation of the expression of the immediate early genes c-fos and egr-1 and in-situ hybridization to count egr1-positive cells we found a specific and selective activation of the caudal part of the dorso-central (Dc) division of the zebrafish pallium upon change in numerosity. As pallial regions are implicated in number cognition in mammals and birds, these findings support the existence of an evolutionarily conserved mechanism for approximate magnitude and provide an avenue for exploring the underlying molecular correlates.

## Introduction

What underlies the ability to deal with numbers and where did it come from? It has been argued that our ability to accurately represent the number of objects in a set (numerosity), and to carry out numerical comparisons and arithmetic, developed from an evolutionarily conserved system for approximating numerical magnitude, the so-called Approximate Number System (ANS, Dehaene, 1997; Feigenson, Dehaene & Spelke, 2004; Gallistel & Gelman, 2000).

The cellular processes and neurocircuitry underlying the operating of the ANS remain to be fully defined; however, subregions of the parietal and prefrontal cortex of human and non-human primates have been identified as plausible candidates (Piazza & Eger 2016; Nieder, 2016; Viswanathan & Nieder, 2013; 2020). In non-human primates, single cell recordings identified neurons that exhibit the expected ANS response, with a peak of activity to one quantity and a progressive drop off in activity as the quantity becomes more distant from the preferred one, in a way that obeys Weber’s Law (Nieder & Miller, 2003; Nieder & Merten, 2007). Similar to the “number neurons” that can be detected in the prefrontal cortex and the ventral intraparietal area in monkeys’ brains, neurons with ANS responses have been identified in crows (Ditz & Nieder, 2015; 2016), within the nidopallium caudolaterale, a brain region that has been argued to be equivalent, though likely not homologous, to the mammalian prefrontal cortex.

A variety of studies have documented non-symbolic numerical competence in a variety of other vertebrate species ranging from non-primate mammals (Utrata, Virányi & Range, 2012; Bánszegi et al., 2016; Perdue et al., 2012; Abramson et al., 2013) to several species of birds (Pepperberg, 2006; Rugani et al., 2009; Rugani et al., 2013; Ditz & Nieder, 2016; see for general reviews Butterworth et al, 2018; Ferrigno & Cantlon, 2017 and referecences therein, and Nieder, 2019; Vallortigara, 2014; 2017). Note that mammals possess a laminated cortex and birds have been shown to possess in their non-laminated pallium circuits organized in lamina-like and column-like entities (Stacho et al., 2020). However, other animals that lack a laminated cortex, such as amphibians (Krusche et al., 2010, Stancher et al., 2015), reptiles (Gazzola et al., 2018; Miletto Petrazzini et al., 2018) and fish (Stancher et al., 2013; Agrillo et al., 2017), show numerical abilities. Interestingly, in all these taxonomic groups (i.e. fish, amphibians, reptiles, birds and mammals) numerosity discrimination exhibits a ratio-dependent signature, in accordance with the Weber law. This might suggest some deep homology in the underlying genetic mechanisms or maybe evolutionary convergence. In order to test the hypothesis of a conserved ANS, a mechanistic, bottom-up approach is needed, with a focus on exploring the neural underpinnings of cognitive features of numerosity, and the genes that control them. Use of zebrafish could be key to such a research, for in recent years it has become established as a developmental and behavioral genetic model species.

Zebrafish have been successfully used for comparative studies of numerosity using conditioning (Potrich et al., 2019; Agrillo et al., 2017), free choice (Pritchard et al., 2001; Potrich et al., 2015; Seguin & Gerlai, 2017) and habituation/dishabituation (Messina et al., 2020) experiments. Variation in the expression of specific immediate early genes (IEGs; Sumbre et al., 2014; Lau et al., 2011) associated with dishabituation to visual numerosity in the telencephalon of zebrafish has been reported (Messina et al., 2020). Here, we refine and extend such analyses to explore the specific nuclei involved in numerical discrimination in the telencephalon of zebrafish.

The zebrafish telencephalon is composed by two main regions: a dorsal region, called pallium, and a ventral region, named subpallium (Northcutt, 1981; Northcutt, 1995; Nieuwenhuys & Meek, 1990). These macroscopical subdivisions can be subdivided into several pallial regions, including the central part of the *area dorsalis telencephali* (Dc), the medial part of the *area dorsalis telencephali* (Dm), the lateral part of the *area dorsalis telencephali* (Dl) (Nieuwenhuys, 2009; Ganz et al., 2015), and into subpallial nuclei such as the *area ventralis telencephali* (V) (Ganz et al., 2012). Each has specific molecular signatures.

In our study, zebrafish were first presented (habituation) with a set of elements (small dots) that changed in individual size, position and density from trial to trial, but remained constant in their numerousness and in the overall areas subtended by the stimuli. Then, a novel visual stimulus was shown (dishabituation) involving controlled changes in different groups of animals: in numerosity, in shape or in size. In a control group, the stimulus remained unchanged. Zebrafish were then sacrificed, their brains were dissected in Dc, Dm, Dl and V, and processed for quantitative polymerase chain reaction (qPCR) analyses of the expression of *c-fos* and *egr-1*. The results were validated by subsequent *in-situ* hybridization assays.

## Materials and Methods

### Ethical Regulations

Experimental procedures complied with the European Legislation for the Protection of Animals used for Scientific Purposes (Directive 2010/63/EU) and were approved by the Scientific Committee on Animal Health and Animal Welfare (Organismo Preposto al Benessere Animale, OPBA) of the University of Trento and by the Italian Ministry of Health (Protocol n. 893/2018-PR and Protocol n. 135/2020-PR).

### Animals

Two hundred and fifty wild-type mixed-strain male nine-month-old zebrafish were used for the behavioural procedures. Eighty of them were randomly selected for qPCR experiments and 80 for *in-situ* hybridization assays. Zebrafish were housed in 7-litre plastic tanks in an automated aquarium system (ZebTEC Benchtop, Tecniplast) and kept separated in groups of 10 individuals based on sex. They were reared in standard conditions (28°C, light/dark cycle of 12h/12h); feeding was provided three times per day using dry food in accordance with guidelines.

### Habituation-Dishabituation experiment

#### Apparatus and stimuli

The setup was the same as in Messina et al. (2020), and consisted of a white plastic arena (40 x 60 x 30 cm) inside of which were placed 5 rectangular smaller tanks (20 x 6.5 x 20 cm, see Fig. 1A) raised 15 cm from the base of the arena, each one housing a single animal. The tanks were made of a white plastic material (Poliplak©) on the four sides, with a white mesh (grid 0.1 mm thick) forming the base, allowing for good water circulation. The water in each tank (8 cm in height) was maintained at a constant temperature of 26°C and kept clean by a pump and a filter system (Micro Jet Filter MCF 40). The apparatus was lit by 2 led strips and a webcam (Microsoft LifeCam Studio) recorded fish behaviour from above (50 cm) the setup.

**Figure 1.**
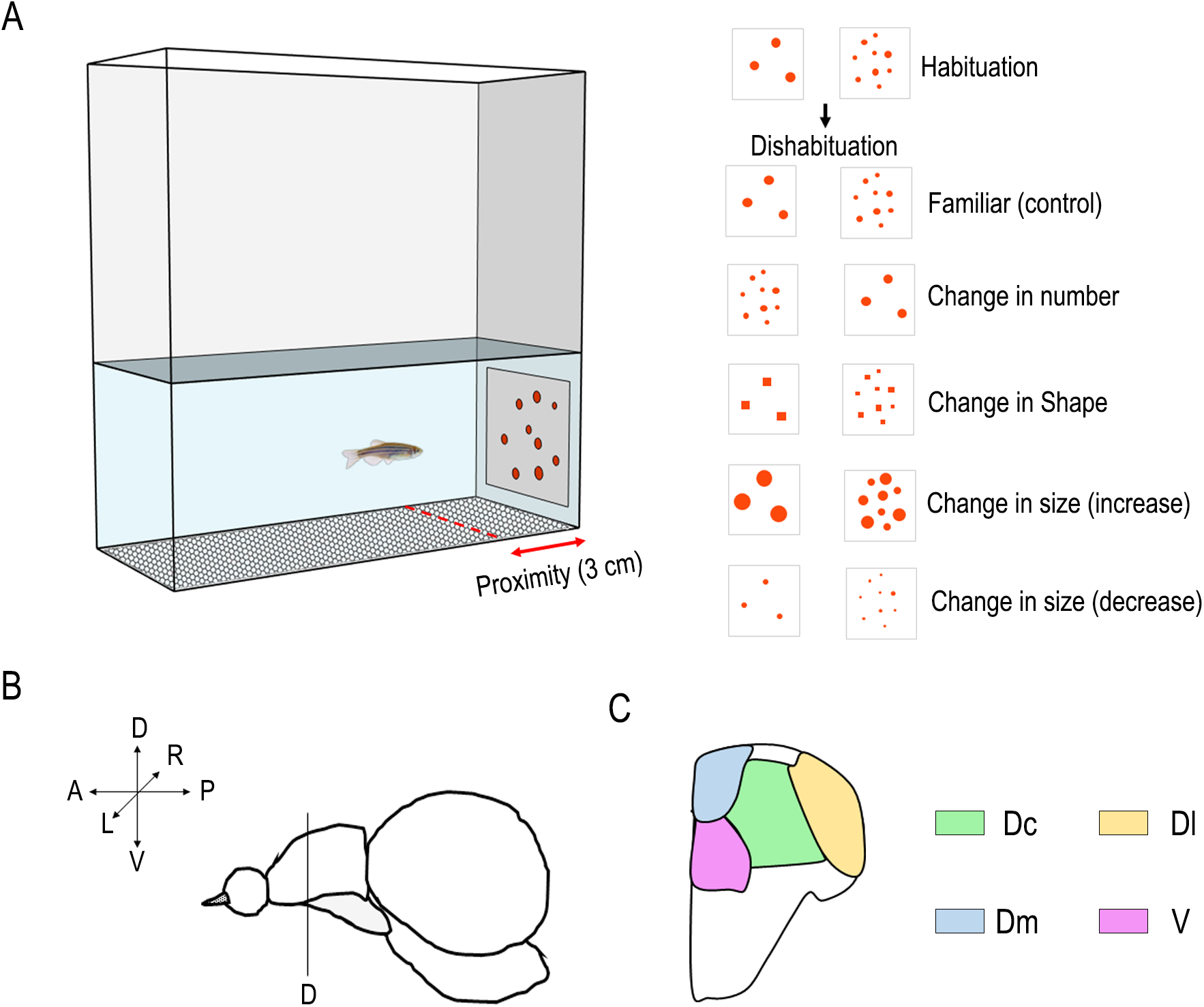
Experimental Design. (A) Apparatus and stimuli used for the habituation and dishabituation phases. Scheme of the lateral view (B) of zebrafish telencephalon with a cross-section of telencephalic nuclei (C) tested for molecular biology analyses. Dc (dorsalcentral), Dl (dorsal-lateral), Dm (dorsal-medial), and V (subpallium).

The stimuli (Fig. 1A) used for the habituation and dishabituation phases were cards (6 × 6 cm) glued on white plastic panels (20 × 6 cm). For the habituation phase, each stimulus depicted a group of 3 or 9 red/orange (RGB: 252,72,11) dots on a white background. For each numerosity, a set of 9 stimuli configurations was used. Among the different configurations, the spatial dispositions of the dots and the size of each dot (range 4-11 mm) were randomized. The overall cumulative area of the stimuli (sum of the dots’ areas) was equalized (1.58 cm^2^) among the different stimuli configurations and the two different numerosities.

For the dishabituation phase, new sets of nine stimuli were used. The novel stimuli comprised: a change in number (from 3 to 9 dots or *vice versa*) keeping the same overall surface area; a change in shape (from dots to squares) maintaining the number and the overall area unmodified; a change in size (increasing or decreasing three times the overall dots surface area) keeping the shape and number unmodified. In each dishabituation stimulus, the spatial distribution of the dots was randomly changed so as to modify continuously density and convex hull, as well as the size of each single element.

### Procedure

Two days before the starting of the experiment, the fish were singly inserted in the apparatus tank in order to acclimatize the animals to the novel environment and reduce the stress connected to isolation. Fish remained in the tank for the entire duration of the experiment, which lasted five days. During the habituation phase, at the beginning of each trial, one panel depicting 3 or 9 dots (depending on the habituation condition) was introduced in one of the two shortest sides of the tank, followed by the release of a small morsel of food (1-1.2 mm) in proximity to the stimulus (after a delay of 30 seconds). The stimulus remained in the tank for 2 minutes after the food delivery and then it was removed. After an inter-trial time of 5 minutes, a new trial started on the opposite side of the tank with a new panel depicting a different dot configuration (but with the same numerosity). Each fish received twelve daily trials, divided in three sessions of four trials each. Among the twelve trials, the configuration of the habituation stimuli was randomized. On the fifth day, fish performed only the first habituation session (four trials). After that, fish were left in their tanks for five hours before the dishabituation test. This delay was to allow the IEG expression to return to the baseline level before the test.

The dishabituation phase consisted of a single trial in which a novel test stimulus was presented to the fish. Before the test, fish were randomly assigned to the five different dishabituation groups that included a change of numerosity (from 3 to 9 dots or *vice versa*, but the same overall area), a change in shape (from dots to squares, but with the same number and overall area), two changes of areas (increasing or decreasing the dots’ surface area, but depicting the same number and shape) or a control condition (same stimulus as used in the habituation phase). In the test trial, the panel was introduced along one of the shortest sides and remained in the tank for 5 minutes. No food was provided during this test trial. Fish were then sacrificed 30 minutes after the end of the dishabituation test and their brains were collected.

As a behavioural measure, we analysed the time spent in proximity of the stimulus (3 cm area) in the 30 seconds after the stimulus appearance. An absolute proportion of time was calculated comparing the dishabituation trial (*test time*) with the previous habituation session (average of the four trials, *habituation time*) performed on the same day, using the following formula:

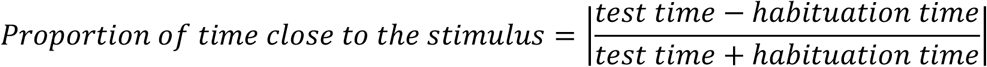

The use of an absolute value for the proportion of time allowed us to detect a behavioural difference between the dishabituation and habituation phases irrespective of whether fish tended to approach or to avoid the novel stimulus compared to the familiar one.

### Tissue preparation: brain dissection and total RNA extraction

Thirty minutes after the end of the dishabituation phase, fish were sacrificed in a bath of ice-cold phosphate-buffered saline solution (PBS; Fisher Bioreagents, USA); their brains were dissected and embedded for later cryosectioning in optimum cutting temperature (OCT, Tissue-Tek OCT Sakura; Sakura Finetek, USA), frozen, and stored at - 20°C. 50 μm coronal sections of the brains were prepared using a cryostat (Leica CM 1860 UV; Leica Biosystems, Germany). Each section was uncurled with fine brushes, put onto a glass slide (Super-Frost Plus; ThermoFisher Scientific, USA) and stored at −20°C. Selected brain areas (central part of the *area dorsalis telencephali* (Dc), medial part of the *area dorsalis telencephali* (Dm), lateral part of the *area dorsalis telencephali* (Dl) and the area *ventralis telencephali* (V); Figure 1B and 1C) were punched out (Li et al., 2018) using 10 μl pipette tips and their total RNA was extracted (Arcturus™ Picopure™ RNA isolation Kit; ThermoFisher Scientific, USA), according to manufacturer’s instructions. Finally, the purity (A260/A280 and A260/230 values) and the concentration of collected total RNAs were assessed using the Nanodrop™ spectrophotometer (Nanodrop™ OneC; ThermoFisher Scientific, USA). Reverse transcription was performed using the SuperScriptTM VILO™ cDNA Synthesis Kit (Invitrogen, ThermoFisher Scientific, USA) according to manufacturer’s instructions.

### Quantitative Polymerase Chain Reaction (qPCR)

Quantitative PCR (qPCR) experiments were performed in order to analyse the expression of *c-fos* (NM_205569), *egr-1* (NM_131248) and *18S* ribosomal RNA (18S) (NM_173234) - which was used as reference gene - and of the molecular markers *emx2, emx3, prox1, eomesa, dlx2a, dlx5a*. Specific primer pairs were commercially synthesized (Sigma-Aldrich/Merck, Germany; see Table 1). qPCR assays were performed in triplicate reactions using the PowerUp™ SYBR™ Green Master Mix (2X) and run in a CFX96™ Real-Time System (Bio-Rad, USA). The ΔCq method was used for expression quantification (Messina et al., 2020). Data were normalized on the expression of the *18S* reference gene (ΔCq) and the relative expression (to the reference gene) of each target was calculated.

**Table 1.**
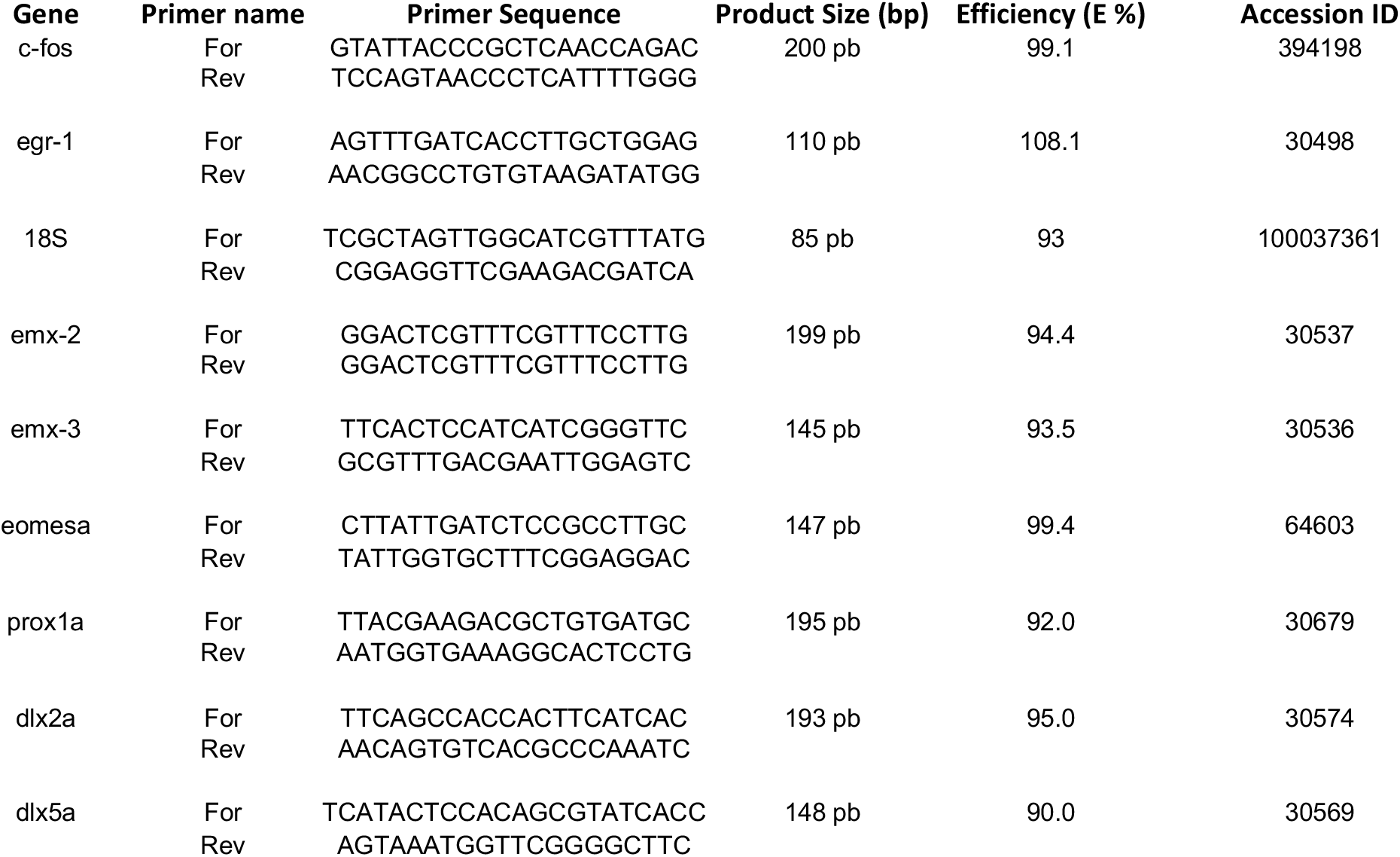
Primers used for qPCR experiments. For: forward primer, Rev: reverse primer.

### In situ hybridization

In-situ hybridization assays were performed to determine the localization of the expression of *egr-1* in the zebrafish brain nuclei. RNA probes necessary for the detection of the *egr-1* mRNA transcripts were created from total brain cDNA by PCR amplification (using the Phusion™ High-Fidelity PCR Master Mix with HF Buffer; ThermoFisher Scientific, USA), followed by precipitation and quantification (Nanodrop™ OneC; ThermoFisher Scientific, USA). Primers for cDNA amplification were as follows: SP6-*egr1*-forward ATTTAGGTGACACTATAGTCTGTTCAGCCTGGTGAGTG, T7-egr1-reverse TAATACGACTCACTATAGTGGAGACCGGAGAAGGGTAAG. DIG-labeled (Digoxigenin-11-UTP/DIG RNA Labeling Mix, Merk, Germany) single-stranded RNA probes were prepared following standard protocols.

The 20 μm brain slices were fixed in 4% paraformaldehyde (Carlo Erba Reagents, Italy), rinsed in PBS and hybridized with *egr1* probes in a humidified chamber at 65° overnight. Then, slides were washed in formamide/SSC solution (formamide: Invitrogen, ThermoFisher Scientific, USA; 20x SSC, saline-sodium citrate buffer: Gibco, ThermoFisher Scientific, USA) at 65°C and in MAB solution (maleic acid buffer; Sigma-Aldrich/Merck, Germany) at room temperature. After being treated with a blocking solution (composed of Fetal Bovine Serum (Euroclone, Italy), Blocking Reagent (Roche, Switzerland), MAB), glass slides were incubated with an anti-DIG-AP antibody (Anti-Digoxigenin-AP, Fab fragments; Merck, Germany) overnight in a humidified chamber. Slides were treated with BCIP/NBT substrate of alkaline phosphatase (BCIP/NBT Ready-To-Use Substrate; SERVA, Germany) and kept in the dark until the colorimetric reaction reached the expected point. Finally, the slides were mounted using Fluoroschield™ with DAPI (Sigma-Aldrich/Merck, Germany) and analysed under a microscope (Observer.Z1, ZEISS, Germany) using a 20x objective and a digital camera (Zeiss AxioCam MRc 5).

Using a single-blind procedure (the operator did not know what training the fish underwent), we counted *egr*-1-positive cells sampling three different rostro-caudal region of Dc according to section 60 (rostral, Dc1), section 85 (medial, Dc2) and section 98 (caudal, Dc3) of a topological atlas of the neuroanatomy of the zebrafish brain (Rupp et al., 1996). Estimated density was reported as number of counted *egr*-1-positive cells (darkblue dots) normalized on the surface of the relative Dc-counted regions in each slice. We used the ZEN Imaging software (Zeiss) for the counting of cells. *egr1*-positive cells were digitally marked using the event marker of the ZEN software, which then provided the total number of positive cells as output.

### Statistical Analysis

Statistical analyses on behaviour, qPCR and *egr1*-positive cells count data were performed using the Statistical Package for the Social Sciences (IBM SPSS Statistics; IBM, USA).

On the behavioural data, an arcsin transformation was used, as recommend for data represented as proportions. An analysis of variance (ANOVA) was performed with “habituation” and “test” as between-subjects factors.

Data for qPCR were analysed with a two-way analyses of variance (applying the Greenhouse-Geisser correction to adjust for the lack of sphericity) using habituation (habituation with either 3 or 9 dots) and type of test [familiar (control condition, no change with respect to the habituation phase), number, shape, surface area increase and surface area decrease] as between-subject factors, and telencephalic nuclei (Dc, Dl, Dm, V) as a within-subject factor. LSD post hoc tests with Bonferroni correction for multiple comparisons were used for pairwise comparisons.

Data for *egr1*-positive cells count were acquired by in situ hybridization using twoway analyses of variance comparing and applying LSD post hoc tests with Bonferroni corrections for multiple comparisons.

## Results

### Behaviour

Proportions of time spent close to the familiar or changed (dishabituated) stimulus is shown in Fig. 2. The Analysis of variance with “habituation” (3 or 9 elements) and “test” (no change (familiar, control group), change in number, change in shape, change in surface area (increase), change in area (decrease)) revealed a significance main effect of the test (F(4,240)= 2.880, p=0.023) but not of the habituation (F(1,240)=0.477, p=0.490) and of the interaction between habituation and test (F(4,240)=0.070, p=0.991). An ANOVA limited to the conditions with a change at test (in number, in shape and in surface areas) did not reveal any statistically significant heterogeneity among conditions (test: F(3,192)=0.523, p=0.667; habituation: F=(1,194)=0.195, p=0.659; habituation x test: F(3,192)=0.049, p= 0.986). Significant differences when the familiar (no change) condition was compared with that of the change in number (t(98)=-2.766, p=0.007), and change in areas (increase, t(98)=-2.901, p=0.005; decrease, t(98)=-3.386, p=0.001), were observed, whereas with the change in shape there was a marginally non-significant effect (t(98)=-1.888, p=0.062).

**Figure 2.**
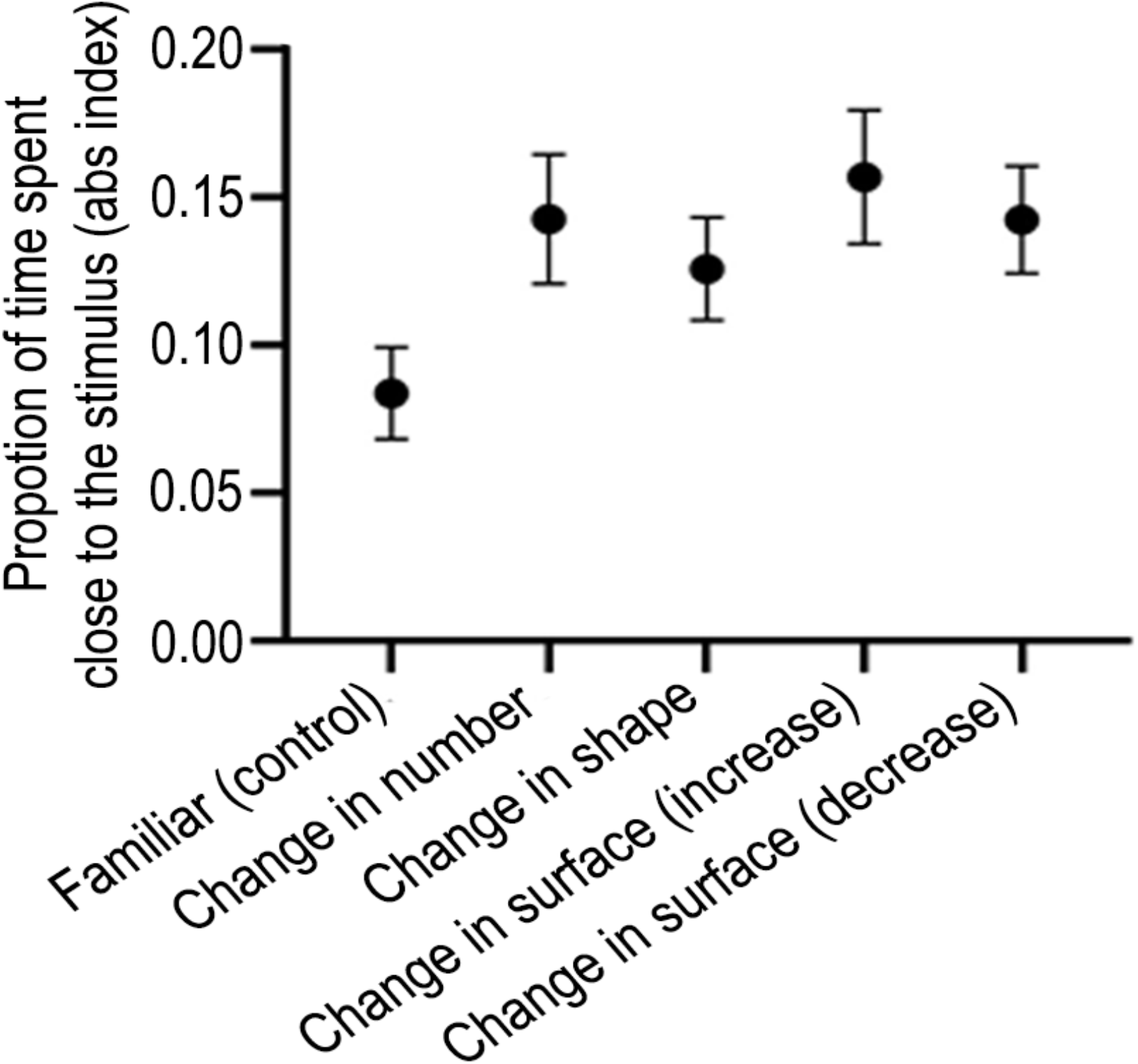
Results of the dishabituation test expressed as the absolute proportion of time spent near the stimulus. Group means with SEM are shown.

Results are shown in Figure 2 (collapsed for the two habituation conditions, i.e. habituation with 3 and 9 dots, since no significant difference between the two types of habituation was observed; separate graphs for the two conditions are however shown in the Supplementary Materials Figure 1).

### Molecular signature analyses for Dc, Dl, Dm and V

In order to assess whether the dissection of the telencephalic nuclei of interest was effective, the expressions of molecular signatures specific for Dc, Dl, Dm and V were measured. The nuclei under investigation are characterized by the expression of some molecular markers (Ganz et al., 2012; Ganz et al., 2015). As reported in the literature we found that in our samples (see Fig. 3) Dc was primarily characterized by the expression of *emx2, emx3* and *eomesa*; Dl by the expression of *emx3, prox1* and *eomesa*; whereas *emx3* alone, was highly expressed in Dm; V was characterized by the expression of *dlx2a* and *dlx5a*, with low *eomesa* mRNA levels (Fig. 3).

**Figure 3.**
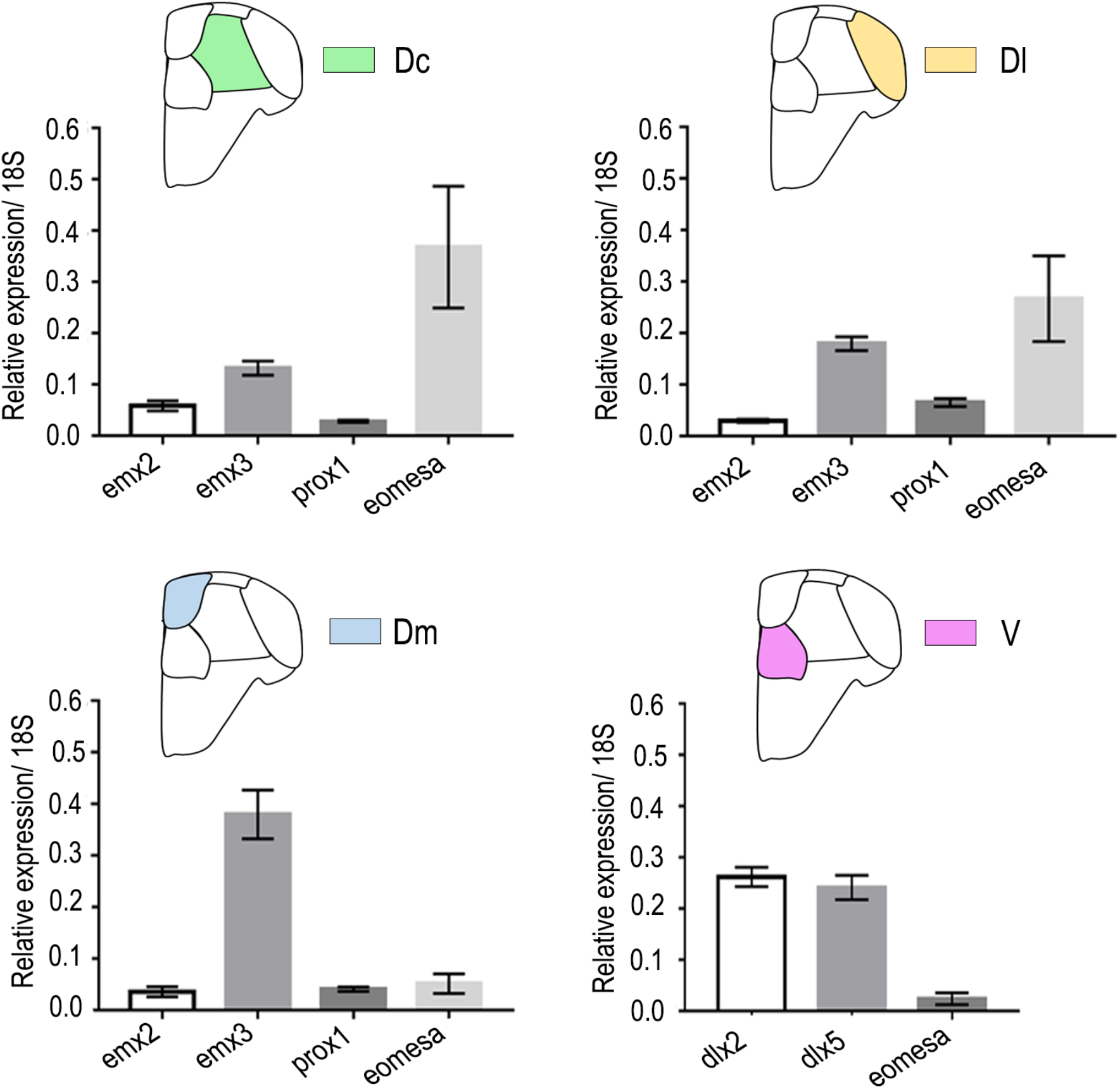
qPCR results for the relative expression of molecular markers in the central part of *area dorsalis telencephali* (A), in the lateral part of *area dorsalis telencephali* (B), in the medial part of *area dorsalis telencephali* (C), and in the *ventral subpallium* (D) in the different test conditions. Group means with SEM are shown. ANOVA revealed a significant main effect of the nuclei for each gene analysed: emx2 (F(2, 192)=3.543, p=0.034; LSD post hoc tests: Dc vs. Dl p=0.015; Dc vs. Dm p=0.043); emx3, (F(2, 192)=19.947, p=0.0001; LSD post hoc tests: Dc vs. Dm p=0.0001); Dl vs. Dm p=0.0001); prox1, (F(2, 192)=12.849, p=0.0001: LSD post hoc tests: Dc vs. Dl p=0.0001; Dl vs. Dm p=0.001); eomesa (F(2, 192)=3.669, p=0.027; LSD post hoc tests; Dc vs. Dm p=0.009).

### Immediate early gene (IEG) expression

Since *c-fos* and *egr-1* are characterized by distinct expression pathways, separate analyses of variance (ANOVA) were performed for the two IEGs, with habituation (habituation with either 3 or 9 dots) and type of test [familiar (control condition, no change with respect to the habituation phase), number, shape, surface area increase and surface area decrease] as between-subject factors, and telencephalic nuclei (Dc, Dl, Dm, V) as a within-subject factor.

Since the overall ANOVA revealed a main effect of the test (*F*(4, 70) = 5.646, *p* = 0.001) and an interaction between telencephalic nuclei and habituation (*F*(2.555, 178.880) = 2.918, *p* = 0.044) for *c-fos*, and a main effect of the telencephalic nuclei (*F*(2.705, 189.319) = 22.083, *p* < 0.0001) and an interaction between telencephalic nuclei and test (*F*(10.818, 189.319) = 2.307, *p* = 0.012) for *egr-1*, in the subsequent analyses we considered test and habituation separately for the distinct telencephalic nuclei (see Supplementary materials for the complete ANOVAs).

### Central part of the area dorsalis telencephali (Dc)

For *c-fos* (see Fig. 4 leftmost column) a comparison between familiar (no change) and change in numerosity revealed a significant test x habituation interaction (F(1, 28) = 25.789, p = 0.0001). Change in numerosity from 3 to 9 resulted in an increase in *c-fos* expression (p=0.0001), whereas change from 9 to 3 in a decrease (p=0.005).

**Figure 4.**
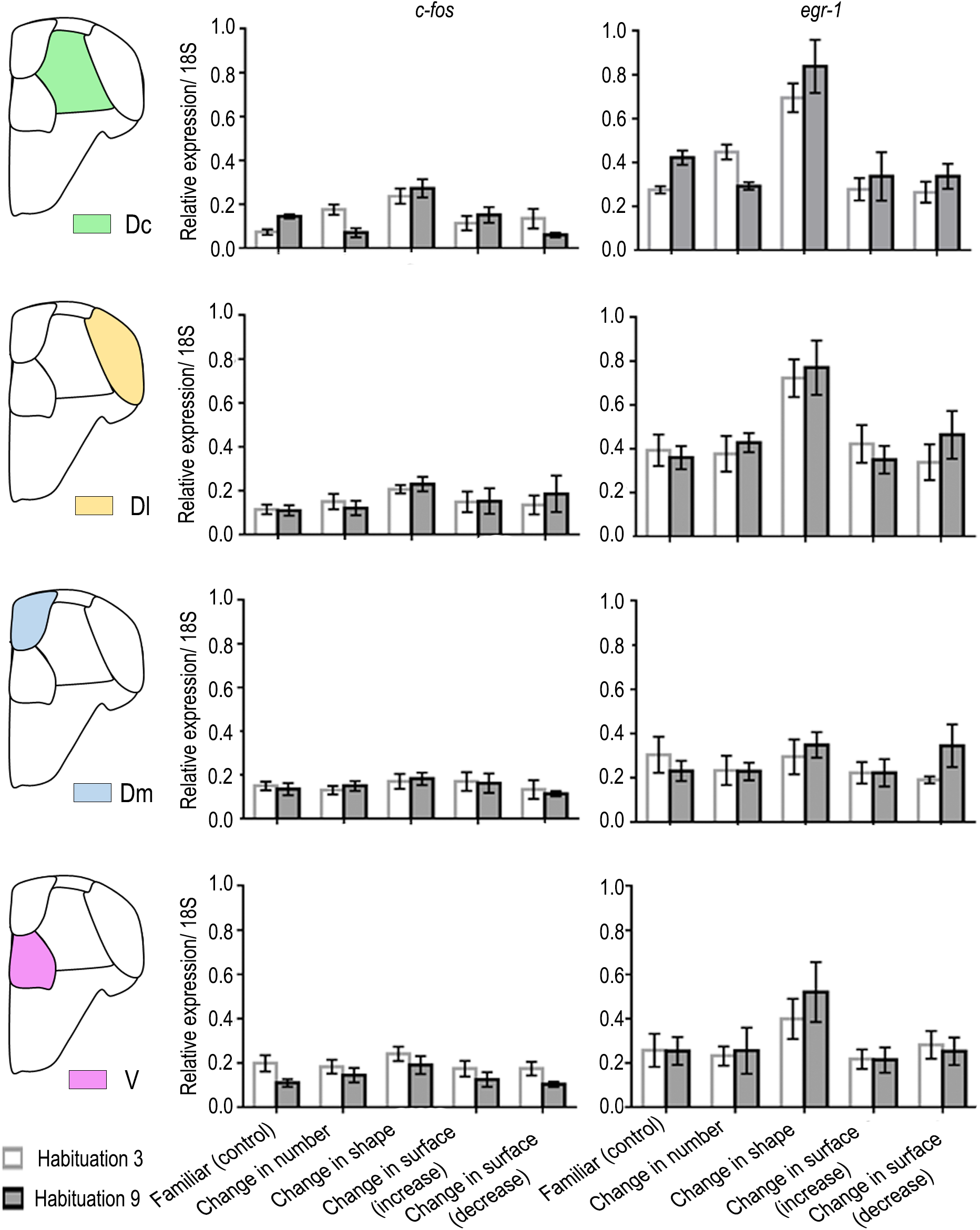
qPCR results for the relative expression of c-fos and egr-1 in the central part of *area dorsalis telencephali* (Dc), in the lateral part of *area dorsalis telencephali* (Dl), in the medial part of *area dorsalis telencephali* (Dm), and in the *ventral subpallium* (V) for the different test conditions. Group means with SEM are shown.

A comparison between familiar (no change) and change in shape revealed only a main effect of the test (F(1, 28) = 26.417, p = 0.0001), with a general increase in *c-fos* expression as a result of the change in shape.

A comparison between familiar (no change) and change in surface area did not reveal any significant main effect of the test (F(2, 42)=0.823, p=0.446) but there was a significant test x habituation interaction (F(2, 42)=3.717, p=0.033). The interaction, however, was limited to the decrease in surface area condition (F(1, 28)=9.077, p=0.005). For egr-1 (Fig. 4 rightmost column), a comparison between familiar (no change) and change in numerosity revealed a significant test x habituation interaction (F(1, 28) = 35.905, p = 0.0001). Similarly to *c-fos*, change in numerosity from 3 to 9 resulted in an increase in egr-1 expression (p=0.0001) whereas change from 9 to 3 in a decrease (p=0.001).

A comparison between familiar (no change) and change in shape revealed only a main effect of the test (F(1, 28) = 35.219, p = 0.0001), with an increase in *erg-1* expression irrespective of habituation with 3 or 9 elements.

A comparison between familiar (no change) and change in size did not reveal any significant main effect or interaction.

Overall, the results suggested that the central part of the *area dorsalis telencephali* (Dc) responded to change in numerosity and shape.

### Lateral part of the area dorsalis telencephali (Dl)

For *c-fos* (Fig 4 rightmost column) no significant effects were observed, whereas only a significant main effect of test was observed for *egr-1* (F(4, 70) = 6.531, p = 0.0001) clearly due to the change in shape.

The results suggested that the lateral part (Dl) of the zebrafish dorsal pallium was not involved in quantity estimation (number and size), but only (though limited to egr-1 expression) in detection of change in shape.

### Medial part of the area dorsalis telencephali (Dm)

No significant main effects or significant interactions were apparent for either of the two IEGs (Fig. 4). The results suggested that the medial part (Dm) of the zebrafish dorsal telencephalon did not show any relevant regulation of neural activity following the different types of changes.

### Area ventralis telencephali (V)

The ANOVA for *c-fos* revealed only a significant main effect of habituation (F(1, 70) = 10.052, p = 0.002). A significant main effect of test was detected for both *c-fos* (F(4, 70) = 3.159, p = 0.019) and *egr-1* (F(4, 70) = 5.487, p = 0.001), limited to the change in shape (Fig. 4). The results suggested that the area ventralis telencephali (V) was not involved in quantity estimation in zebrafish, either discrete (numerosity) or continuous (surface area) but only in shape.

### Counting of egr-1-positive cells

Real time qPCR showed that Dc was the only area that showed modulation of expression of both *c-fos* and *egr-1* to a change in numerosity. However, it also showed modulation of response to change in shape. We thus looked at the spatial location of *egr*-1-positive cells that respond to numerosity and shape along the rostro-caudal axis of Dc using in situ hybridization (Fig. 5). (We did not show *c-fos* positive cells, due to the weakness of the detected signal in our experiments).

**Figure 5.**
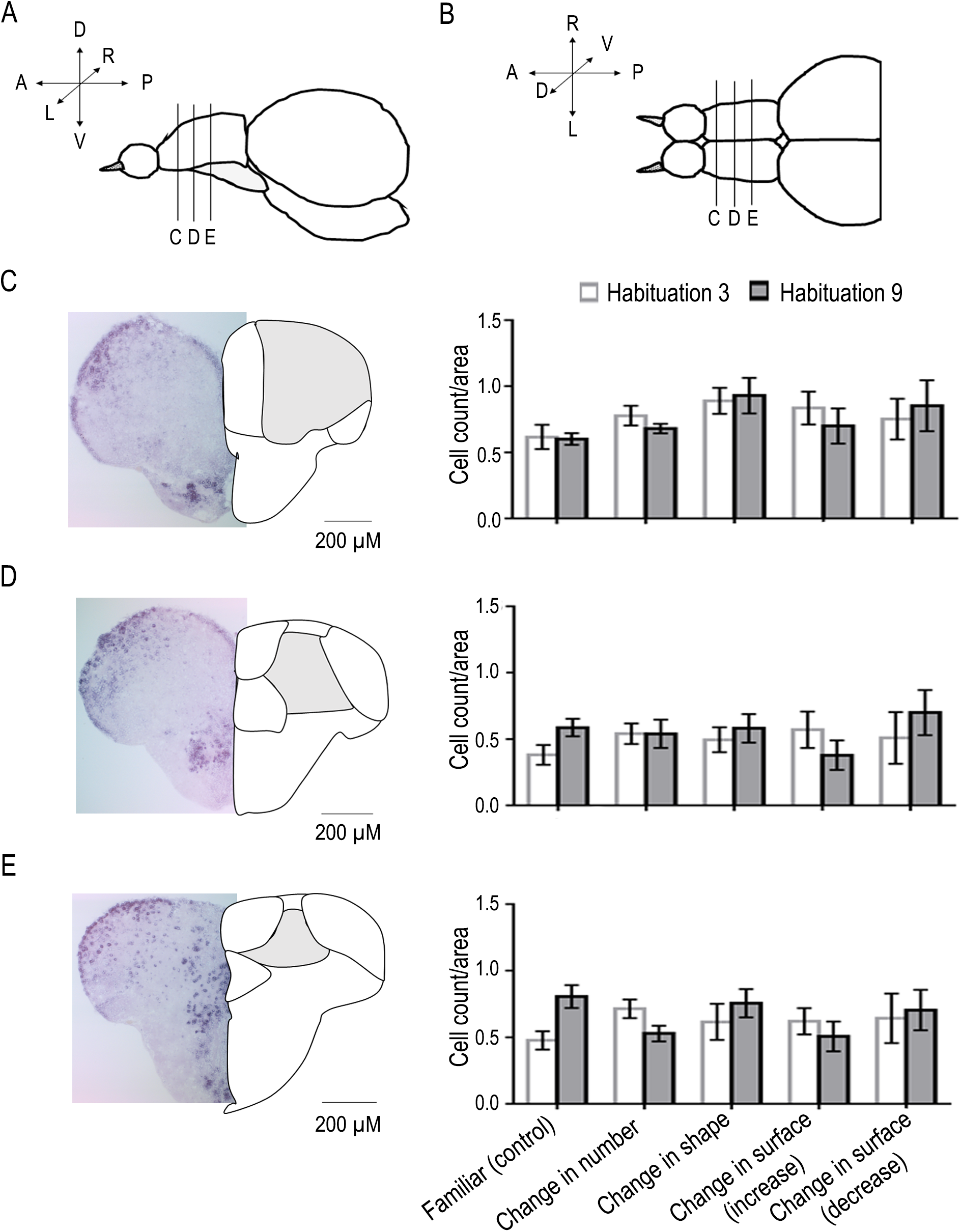
In situ hybridization analysis of *egr-1*. Mean number of *egr-1*-positive cells in three different rostro-caudal regions of Dc. Scheme of lateral (A) and dorsal (B) view of zebrafish telencephalon with results for the selected rostral (C), medial (D) and caudal (E) slices in the different test conditions. (Group means with SEM are shown.)

An ANOVA was run in order to evaluate the percentage of *egr-1*-expressing cells in the three Dc slices along a rostro-caudal position. The ANOVA revealed a significant main effect of the rostro-caudal position of the slice (*F*(2, 140) = 42.360, *p* < 0.0001) and a significant interaction of the rostro-caudal position with test (*F*(8, 140) = 2.168, *p* = 0.033).

In particular, as shown in Fig. 5c, in the most rostral region, Dc1, a comparison between familiar (no change) and change in shape revealed a significant main effect of the test (F(1,28) = 9.422, p = 0.005). No significant main effect nor significant interaction were observed in the medial region of Dc (Dc2, figure 5 D). In the most caudal region of Dc (Dc3, figure 5 E) a significant interaction between habituation and test (F(2, 56)=12.907, p=0.001) was observed, with an increase (p=0.026) or a decrease (p=0.011) in cell count depending on whether the change in the stimulus consisted of an increase or a decrease in numerosity.

Thus, *in situ* hybridization results suggested that the most rostral part of Dc was responsive to a change in shape whereas the most caudal part to a change in numerosity.

## Discussion

The habituation-dishabituation design of the stimulus presentation allows the disentanglement of the effect of changing stimulus numerosity, stimulus shape and stimulus size. The results of qPCR experiments showed that different regions of the zebrafish telencephalon differentially expressed *c-fos* and *egr-1* depending on the kind of change in the stimulus. The central part of the area dorsalis telencephali (Dc), the lateral part of the area dorsalis telencephali (Dl), and the area ventralis telencephali (V) were all affected by changes in shape. Note, however, that changes in shape likely represented different aspects of the stimulus, such as shape in itself and spatial aspects. The DI area in teleosts has been proposed to be homologous to the hippocampal formation of mammals and birds (Rodríguez et al., 2002; Teles et al., 2015); thus, perhaps the modulation of response to shape in DI reflects selectivity to change in the spatial characteristics of the stimulus (note that in this area, selectivity in expression to change in shape was apparent only for *egr-1* but not for *c-fos*).

Selectivity of response to numerosity was confined to Dc only. This area also responded to shape but in situ hybridisation showed that the rostral part only responded to shape whereas the most caudal part only responded to numerosity. More precisely, we found that a larger number of *egr-1* - expressing neurons was seen in fish habituated with 3 dots and tested with 9, and a smaller number in those habituated with 9 dots and tested with 3, suggesting that the increased or decreased expression of *egr-1* mRNA in qPCR experiments was probably due to a larger or smaller number of activated neurons recruited during the dishabituation phase in fish facing the numerosity change. This pattern of response is reminiscent of properties of neurons in the lateral intraparietal area of monkeys’ brain that showed increased or decreased activity as a function of the number of elements entering their receptive fields, thus encoding the number of elements in a visual array in a monotonic manner (Roitman et al., 2007).

Selectivity of response to change in surface areas were also limited to only Dc area, however it appeared to be quite small and variable, depending on whether the change involved an increase or a decrease and whether it applied to large (9) or small (3) numerosities and then only for *c-fos* and not for *egr-1*. Considering that some theoretical accounts of number cognition assume that dealing with discrete (countable) numerosities is one aspect of a more general system dealing with magnitude (either discrete or continuous (see e.g.: Gallistel, 1989; Walsh, 2003; and for empirical evidence e.g.: De Corte et al, 2017; Bortot et al., 2020) one would expect clear and parallel responsivity to changes in discrete (numerosities) and continuous (surface area) quantities. It could be, however, that the lack of control for distance of seeing made absolute size estimation difficult for zebrafish. Alternatively, it may be that processing for continuous magnitude is done mainly at the level of the tectum (which has been shown to be responsive for changes in surface area in habituation/dishabituation experiments, Messina et al., 2020) and that the telencephalon is mainly involved with discrete quantities.

In the mammalian brain the main area involved in numerosity cognition is the posterior part of the parietal cortex (Piazza & Eger 2016; Viswanathan & Nieder, 2013, 2020; Nieder, 2016). Activation of the prefrontal cortex is also observed but in single cell recording experiments it usually occurs with a latency of about 30 ms, suggesting a later stage of processing (Viswanathan & Nieder, 2013). In the avian brain only single cell recording experiments are available as of yet, and they suggest that the nidopallium caudolaterale (NCL) in crows contain number neurons similar to those recorded in the mammalian/prefrontal cortex (Ditz & Nieder, 2015; 2016; Nieder, 2016; 2017). The possible homology/homoplasy relationship of NCL with regions of the mammalian brain is at present uncertain. Functionally, NCL appears to be a sort of avian equivalent of the prefrontal cortex (Güntürkün, 2005) but there are also striking differences (e.g., an apparent lack of direct connection between the NCL and the hippocampal formation). Thus, our finding of a highly selective role of the most caudal part of Dc in numerosity responsiveness in zebrafish is exciting in terms of the possible similarities of this region with equivalent or homologous regions in the mammalian and avian brains.

There is some (though admittedly not unanimous) consensus that Dl of teleosts is homologous to the medial pallium of tetrapods (i.e. hippocampal formation), Dm to ventral pallium (pallial amygdala), and Dc to dorsal pallium (note that the mammalian isocortex is one example of the many outcomes of the evolution of vertebrate dorsal pallium, Tosches & Laurent, 2019). Harvey-Girard et al. (2012) in particular hypothesized a homology of Dc with efferent layers V and VI of mammalian isocortex. No data are, however, currently available to dissect anatomically or functionally different parts of Dc, such as the most caudal and rostral regions.

Taken together, our results offer evidence that the central part of *area dorsalis telencephali* (Dc) may be the pallial structure of the zebrafish brain most involved in cognitive processes such as shape and numerosity recognition.

## Author Contributions

Conceived and designed the experiments: A.M, D.P., C.H.B., S.E.F., G.V. Performed the experiments: A.M., D.P., I.S. Analysed and interpreted the data: A.M., D.P., I.S., V.A.S., C.H.B., S.E.F., G.V. Contributed reagents/materials: G.V. All authors contributed to the manuscript writing.

## Acknowledgments

We thank Tommaso Pecchia, Grazia Gambardella, Roberta Guidolin and Ciro Petrone for technical and administrative support. The authors would like to thank Jose Torres-Perez and Eva Sheardown for their useful comments and suggestions.

## References

Abramson, J.Z., Hernández-Lloreda, V., Call, J. & Colmenares, F. Relative quantity judgments in the beluga whale (Delphinapterus leucas) and the bottlenose dolphin (Tursiops truncatus). Behav Processes, 96, 11–19 (2013).

Agrillo C, Piffer L, Bisazza A, Butterworth Evidence for Two Numerical Systems That Are Similar in Humans and Guppies. PLoS ONE 7(2): e31923. (2012)

Bánszegi, O., Urrutia, A., Szenczi, P. & Hudson, R. More or less: spontaneous quantity discrimination in the domestic cat. Anim Cogn, 19 (5), 879–88 (2016).

Bortot, M., Stancher, G., Vallortigara, G. Transfer from number to size reveals abstract coding of magnitude in honeybees. iScience, 23 (5), 101122 (2020).

De Corte, B.J., Navarro, V.M., Wasserman, E.A. Non-cortical magnitude coding of space and time by pigeons. Curr Biol, 27 (23), R1264–R1265 (2017).

Dehaene, S. The number sense: How the mind creates mathematics. New York: Oxford University Press (1997).

Ditz, H.M. & Nieder, A. Neurons selective to the number of visual items in the corvid songbird endbrain. Proc Natl Acad Sci, 112, 7827–7832 (2015).

Ditz, H.M. & Nieder, A. Numerosity representations in crows obey the Weber-Fechner law. Proc Biol Sci, 283 (1827), 20160083 (2016).

Ferrigno, S. & Cantlon, J.F. Evolutionary constraints on the emergence of human mathematical concepts. In J. Kaas (Ed.), Evolution of Nervous Systems (2nd ed. vol. 3), pp. 511–521. Oxford: Elsevier (2017).

Feigenson, L., Dehaene, S., & Spelke, E. Core systems of number. Trends Cogn Sci, 8, 307–314 (2004).

Gallistel, C.R. Animal cognition: the representation of space, time and number. Annu Rev Psychol, 40 (1), 155–189 (1989).

Gallistel, C.R., Gelman, I.I. Non-verbal numerical cognition: from reals to integers. Trends in Cognitive Sciences, 4 (2), 59–65 (2000).

Ganz, J., Kaslin, J., Freudenreich, D., Machate, A., Geffarth, M., & Brand, M. Subdivisions of the adult zebrafish subpallium by molecular marker analysis. J Comp Neurol, 520, 633–655 (2012).

Ganz, J., Kroehne, V., Freudenreich, D., Machate, A., Geffarth, M., Braasch, I., Kaslin, J., & Brand, M. Subdivisions of the adult zebrafish pallium based on molecular marker analysis. F1000Research, 3 (2015).

Gazzola, A., Vallortigara, G., & Pelliteri-Rosa, D. Continuous and discrete quantity discrimination in tortoises. Biol Lett, 14 (12), 20180649 (2018).

Güntürkün, O. The avian ‘prefrontal cortex’ and cognition. Curr Opin Neurobiol, 15 (6), 686–693 (2005).

Krusche, P., Uller, C. & Dicke, U. Quantity discrimination in salamanders. J. Exp. Biol. 213(11), 1822–1828 (2010).

Lau, B.Y.B., Mathur, P., Gould, G.G., & Guo, S. Identification of a brain center whose activity discriminates a choice behavior in zebrafish. Proc Natl Acad Sci, 108, 2581–2586 (2011).

Li, C.Y., Hofmann, H.A., Harris, M.L., & Earley, R.L. Real or fake? Natural and artificial social stimuli elicit divergent behavioural and neural responses in mangrove rivulus, *Kryptolebias marmoratus*. Proc Biol Sci, 14, 285 (2018).

Messina, A., Potrich, D., Schiona, I., Sovrano, V.A., Fraser, S.E., Brennan, C.H., & Vallortigara, G. Response to change in the number of visual stimuli in zebrafish: A behavioural and molecular study. Sci Rep, 10, 5769 (2020).

Miletto Petrazzini, M.E., Bertolucci, C. & Foà, A. Quantity discrimination in trained lizards (Podarcis sicula). Front Psychol, 9, 274 (2018).

Nieder, A. & Miller, E. K. Coding of cognitive magnitude: compressed scaling of numerical information in the primate prefrontal cortex. Neuron 37, 149–157 (2003).

Nieder, A. & Merten, K.A. Labeled-Line Code for Small and Large Numerosities in the Monkey Prefrontal Cortex. J Neurosci, 27, 5986–5993 (2007).

Nieder, A. The neuronal code for number. Nat Rev Neurosci, 17 (6), 366–82 (2016).

Nieder, A. Evolution of cognitive and neural solutions enabling numerosity judgements: lessons from primates and corvids. Philos Trans R Soc B Biol Sci, 373, 1740 (2017).

Nieder, A. A Brain for Numbers: The biology of the number instinct. Mit Press, (2019)

Nieuwenhuys, R. & Meek, J. The Telencephalon of Actinopterygian Fishes. In Jones, E.G. & Peters, A. (eds), Comparative Structure and Evolution of Cerebral Cortex, Part I, Cerebral Cortex. Springer US, Boston, MA, 31–73 (1990).

Nieuwenhuys, R. The forebrain of actinopterygians revisited. Brain Behav Evol, 73(4), 229–252 (2009).

Northcutt, R.G. Evolution of the Telencephalon in Nonmammals. Annu Rev Neurosci, 4, 301–350 (1981).

Northcutt, R.G. The forebrain of gnathostomes: in search of a morphotype. Brain Behav Evol, 46, 275–288 (1995).

Pepperberg, I.M. Grey parrot numerical competence: a review. Anim Cogn, 9 (4), 377–391 (2006).

Perdue, B.M., Talbot, C.F., Stone, A.M. & Beran, M.J. Putting the elephant back in the herd: elephant relative quantity judgments match those of other species. Anim Cogn, 15 (5), 955–961 (2012).

Piazza, M. & Eger, E. Neural foundations and functional specificity of number representations. Neuropsychologia, Special Issue: Functional Selectivity in Perceptual and Cognitive Systems - A Tribute to Shlomo Bentin (1946-2012), 83, 257–273 (2016).

Potrich, D., Sovrano, V.A., Stancher, G. & Vallortigara, G. Quantity discrimination by zebrafish (Danio rerio). J Comp Psychol, 129, 388–393 (2015).

Potrich, D., Rugani, R., Sovrano, V.A., Regolin, L. & Vallortigara, G. Use of numerical and spatial information in ordinal counting by zebrafish. Sci Rep, 9, 18323 (2019).

Pritchard, V.L., Lawrence, J., Butlin, R.K. & Krause J. Shoal choice in zebrafish, Danio rerio: the influence of shoal size and activity. Anim Behav, 62, 1085–1088 (2001).

Rodríguez, F., López, J.C., Vargas, J.P., Gómez, Y., Broglio, C. & Salas, C. Conservation of Spatial Memory Function in the Pallial Forebrain of Reptiles and Ray-Finned Fishes. J Neurosci, 22, 2894–2903 (2002).

Roitman, J.D., Brannon, E.M. & Platt, M.L. Monotonic coding of numerosity in macaque lateral intraparietal area. PLoS Biology, e208 (2007).

Rugani, R., Fontanari, L., Simoni, E., Regolin, L. & Vallortigara, G. Arithmetic in newborn chicks. Proc Biol Sci, 276 (1666), 2451–2460 (2009).

Rugani, R., Vallortigara, G. & Regolin, L. Numerical Abstraction in Young Domestic Chicks (Gallus gallus). PLoS ONE, 8 (6), e65262 (2013)

Seguin, D. & Gerlai, R. Zebrafish prefer larger to smaller shoals: analysis of quantity estimation in a genetically tractable model organism. Anim Cogn, 20 (5), 813–821 (2017).

Stacho, M., Herold, C., Rook, N., Wagner, H., Axer, M., Amunts, K. & Güntürkün, O. A cortex-like canonical circuits in the avian forebrain. Science, 369, 1585 (2020).

Stancher, G. et al. Discrimination of small quantities by fish (redtail splitfin, Xenotoca eiseni). Anim. Cogn. 16 (2), 307–12 (2013).

Stancher, G. et al. Numerical discrimination by frogs (Bombina orientalis). Anim. Cogn. 18 (1), 219–229 (2015).

Sumbre, G. & de Polavieja, G. G. The world according to zebrafish: how neural circuits generate behavior. Front Neural Circuits, 8 (91), (2014).

Teles, M.C., Almeida, O. & Oliveira, R.F. Social interactions elicit rapid shifts in functional connectivity in the social decision-making network of zebrafish. Proc Biol Sci, 282, 1816 (2015).

Tosches, M.A. & Laurent, G. Evolution of neuronal identity in the cerebral cortex. Curr Opin Neurobiol, 56, 199–208 (2019).

Utrata, E., Virányi, Z. & Range, F. Quantity Discrimination in Wolves (Canis lupus). Front Psychol, 16 (3), 505 (2012).

Vallortigara, G. An animal’s sense of number. In “The nature and Development of Mathematics. Cross Disciplinary Perspective on Cognition, Learning and Culture’’ (Adams, J.W., Barmby P., Mesoudi, A., eds.), pp. 43–65, Routledge, New York. (2017)

Vallortigara, G. Foundations of Number and Space Representations in Non-Human Species. In “Evolutionary Origins and Early Development of Number Processing’’, pp. 35–66 (Eds., D.C. Geary, D.B. Bearch, K. Mann Koepke), Elsevier, New York. (2014)

Viswanathan, P. & Nieder, A. Neuronal correlates of a visual “sense of number” in primate parietal and prefrontal cortices. Proc Natl Acad Sci USA, 110 (27), 11187–11192 (2013).

Viswanathan, P. & Nieder, A. Spatial Neuronal Integration Supports a Global Representation of Visual Numerosity in Primate Association Cortices. J Cogn Neurosci, 32 (6), 1184–1197 (2020).

Walsh, V. A theory of magnitude: common cortical metrics of time, space and quantity. Trends Cogn Sci, 7 (11), 483–488 (2003).

Rupp, B., Wullimann, M.F. & Reichert, H. The zebrafish brain: a neuroanatomical comparison with the goldfish. Anat Embryol (Berl). 194 (2), 187–203 (1996).

