## Supplementary Materials for "Neurons in the dorso-central division of zebrafish pallium respond to change in visual numerosity"

**Supplementary Materials of "Neurons in the dorso-central part of zebrafish pallium encode visual numerosity"**

Supplementary Materials Figure 1: Proportion of time spent near the stimulus during dishabituation (comparing the dishabituation trial with the first of the four trials previously performed during the last habituation session) as a function of habituation conditions (with 3 or 9 dots) and test conditions [no change (familiar), change in number, change in shape, change in surface area (increase), change in surface area (decrease).

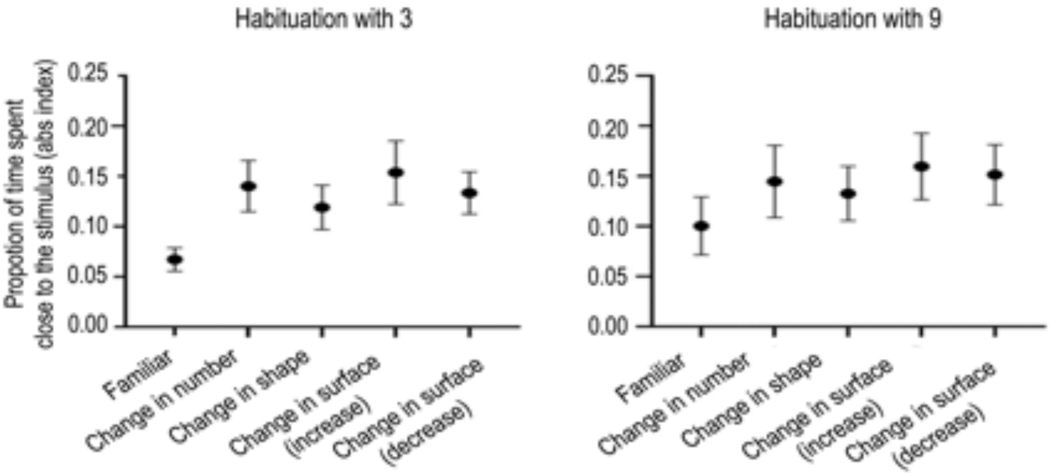

Supplementary Table 1: Overall analyses of variance (ANOVA) for *c-fos* and for *egr-1* in qPCR, with habituation (habituation with 3 dots, habituation with 9 dots) and type of test (familiar, number, shape, surface area increase, surface area decrease) as between-subject factors, and telencephalic nuclei (Dc, Dl, Dm, V) as a within-subject factor.

| ANOVA | | | |
|---|---|---|---|
| c-fos | Main effect of Telencephalic Nuclei | F(2.55,178.880)=0.712 | p=0.524 |
| | Main effect of Habituation | F(1,70)=1.270 | p=0.264 |
| | Main effect of Test | F(4,70)=5.646 | p=0.001 |
| | Telencephalic Nuclei x Test interaction | F(10.222,178.880)=1.444 | P=0.163 |
| | Habituation x Test interaction | F(4,70)=0.437 | p=0.781 |
| | Telencephalic Nuclei x Habituation interaction | F(2.555,178.880)=2.918 | p=0.044 |
| | Telencephalic Nuclei x Test x Habituation interaction | F(10.222,178.880)=1.324 | p=0.219 |
| egr-1 | Main effect of Telencephalic Nuclei | F(2.705,189.3199)=22.083 | p=0.0001 |
| | Main effect of Habituation | F(1,70)=1.241 | p=0.269 |
| | Main effect of Test | F(4,70)=15.217 | p=0.0001 |

| | | | |
|---|---|---|---|
| | Telencephalic Nuclei x Test interaction | F(10.818,189.319)=2.307 | p=0.012 |
| | Habituation x Test interaction | F(4,70)=0.975 | p=0.427 |
| | Telencephalic Nuclei x Habituation interaction | F(2.705,189.319)=0.098 | p=0.950 |
| | Telencephalic Nuclei x Test x Habituation interaction | F(10.818,189.319)=0.853 | p=0.585 |

Supplementary Table 2: Analyses of variance (ANOVA) for *c-fos* and for *egr-1* mRNA expression level in qPCR for Dc, with habituation (habituation with 3 dots, habituation with 9 dots) and type of test (familiar, number, shape, surface area increase, surface area decrease) as between-subject factors.

| Dc | | | |
|---|---|---|---|
| c-fos | Main effect of Habituation | F(1,70)=0.175 | p=0.677 |
| | Main effect of Test | F(4,70)=9.484 | P=0.0001 |
| | Habituation x Test interaction | F(4,70)=3.838 | p=0.007 |
| egr-1 | Main effect of Habituation | F(1,70)=1.156 | p=0.286 |
| | Main effect of Test | F(4,70)=19.382 | p=0.0001 |
| | Habituation x Test interaction | F(4,70)=2.306 | p=0.067 |

Supplementary Table 3: Analyses of variance (ANOVA) for *c-fos* and for *egr-1* mRNA expression level in qPCR for Dl, with habituation (habituation with 3 dots, habituation with 9 dots) and type of test (familiar, number, shape, surface area increase, surface area decrease) as between-subject factors.

| Dl | | | |
|---|---|---|---|
| c-fos | Main effect of Habituation | F(1,70)=0.35 | p=0.852 |
| | Main effect of Test | F(4,70)=1.917 | p=0.117 |
| | Habituation x Test interaction | F(4,70)=0.209 | p=0.933 |
| egr-1 | Main effect of Habituation | F(1,70)=0.168 | p=0.683 |
| | Main effect of Test | F(4,70)=6.531 | p=0.0001 |
| | Habituation x Test interaction | F(4,70)=0.406 | p=0.804 |

Supplementary Table 4: Analyses of variance (ANOVA) for *c-fos* and for *egr-1* mRNA expression level in qPCR for Dm, with habituation (habituation with 3 dots, habituation with

9 dots) and type of test (familiar, number, shape, surface area increase, surface area decrease) as between-subject factors.

| | | | |
|---|---|---|---|
| Dm | | | |
| c-fos | Main effect of Habituation | F(1,70)=0.004 | p=0.949 |
| | Main effect of Test | F(4,70)=0.939 | p=0.447 |
| | Habituation x Test interaction | F(4,70)=0.142 | p=0.966 |
| egr-1 | Main effect of Habituation | F(1,70)=0.189 | p=0.665 |
| | Main effect of Test | F(4,70)=0.938 | p=0.447 |
| | Habituation x Test interaction | F(4,70)=0.578 | p=0.680 |

Supplementary Table 5: Analyses of variance (ANOVA) for *c-fos* and for *egr-1* mRNA expression level in qPCR for V, with habituation (habituation with 3 dots, habituation with 9 dots) and type of test (familiar, number, shape, surface area increase, surface area decrease) as between-subject factors.

| | | | |
|---|---|---|---|
| V | | | |
| c-fos | Main effect of Habituation | F(1,70)=10.052 | p=0.002 |
| | Main effect of Test | F(4,70)=3.159 | p=0.019 |
| | Habituation x Test interaction | F(4,70)=0.282 | p=0.889 |
| egr-1 | Main effect of Habituation | F(1,70)=0.882 | p=0.351 |
| | Main effect of Test | F(4,70)=5.487 | p=0.001 |
| | Habituation x Test interaction | F(4,70)=0.858 | p=0.493 |

Supplementary Table 6: Overall analyses of variance (ANOVA) for *egr-1* positive cells in three different rostro-caudal regions of Dc, with habituation (habituation with 3 dots, habituation with 9 dots) and type of test (familiar, number, shape, surface area increase, surface area decrease) as between-subject factors, and rostro-caudal regions (Dc1, Dc2, Dc3) as a within-subject factor.

| | | | |
|---|---|---|---|
| ANOVA | | | |
| egr-1 | Main effect of rostro-caudal regions of Dc | F(2, 140)=42.360 | p=0.0001 |
| | Main effect of Habituation | F(1,70)=0.163 | p=0.688 |
| | Main effect of Test | F(4,70)=0.569 | p=0.686 |
| | Rostro-caudal regions of Dc x Test interaction | F(8,140)=2.168 | p=0.033 |

| | | | |
|---|---|---|---|
| | Habituation x Test interaction | F(4,70)=0.841 | p=0.504 |
| | Rostro-caudal regions of Dc x Habituation interaction | F(2,140)=1.416 | p=0.033 |
| | Rostro-caudal regions of Dc x Test x Habituation interaction | F(8,140)=1.469 | p=0.174 |

Supplementary Table 7: Analyses of variance (ANOVA) for *egr-1* positive cells in the rostro-caudal regions of Dc1, Dc2 and Dc3, with habituation (habituation with 3 dots, habituation with 9 dots) and type of test (familiar, number) as between-subject factors.

| | | | |
|---|---|---|---|
| Dc1 | | | |
| egr-1 positive cells | Main effect of Habituation | F(1,28)=0.745 | p=0.395 |
| | Main effect of Test | F(1,28)=3.291 | p=0.080 |
| | Habituation x Test interaction | F(1,28)=0.142 | p=0.539 |
| Dc2 | | | |
| egr-1 positive cells | Main effect of Habituation | F(1,28)=1.548 | p=0.224 |
| | Main effect of Test | F(1,28)=0.461 | p=0.503 |
| | Habituation x Test interaction | F(1,28)=1.570 | p=0.221 |
| Dc3 | | | |
| egr-1 positive cells | Main effect of Habituation | F(1,28)=0.990 | p=0.328 |
| | Main effect of Test | F(1,28)=0.077 | p=0.784 |
| | Habituation x Test interaction | F(1,28)=12.907 | p=0.01 |

Supplementary Table 8: Analyses of variance (ANOVA) for *egr-1* positive cells in the rostro-caudal regions of Dc1, Dc2 and Dc3, with habituation (habituation with 3 dots, habituation with 9 dots) and type of test (familiar, shape) as between-subject factors.

| | | | |
|---|---|---|---|
| Dc1 | | | |
| egr-1 positive cells | Main effect of Habituation | F(1,28)=0.014 | p=0.905 |
| | Main effect of Test | F(1,28)=9.422 | p=0.05 |
| | Habituation x Test interaction | F(1,28)=0.079 | p=0.780 |
| Dc2 | | | |
| egr-1 positive cells | Main effect of Habituation | F(1,28)=2.796 | p=0.106 |
| | Main effect of Test | F(1,28)=0.379 | p=0.546 |
| | Habituation x Test interaction | F(1,28)=0.479 | p=0.494 |
| Dc3 | | | |
| egr-1 positive cells | Main effect of Habituation | F(1,28)=5.260 | p=0.030 |
| | Main effect of Test | F(1,28)=0.192 | p=0.665 |
| | Habituation x Test interaction | F(1,28)=0.362 | p=0.362 |

Supplementary Table 8: Analyses of variance (ANOVA) for *egr-1* positive cells in the rostro-caudal regions of Dc1, Dc2 and Dc3, with habituation (habituation with 3 dots, habituation with 9 dots) and type of test (familiar, surface area increase, surface area decrease) as between-subject factors.

| | | | |
|---|---|---|---|
| Dc1 | | | |
| egr-1 positive cells | Main effect of Habituation | F(1,42)=0.024 | p=0.878 |
| | Main effect of Test | F(2,42)=1.216 | p=0.307 |
| | Habituation x Test interaction | F(2,42)=0.399 | p=0.673 |
| Dc2 | | | |
| egr-1 positive cells | Main effect of Habituation | F(1,42)=0.393 | p=0.534 |
| | Main effect of Test | F(2,42)=0.584 | p=0.562 |
| | Habituation x Test interaction | F(2,42)=1.415 | p=0.254 |
| Dc3 | | | |
| egr-1 positive cells | Main effect of Habituation | F(1,42)=0.842 | p=0.364 |
| | Main effect of Test | F(2,42)=0.420 | p=0.660 |
| | Habituation x Test interaction | F(2,42)=1.617 | p=0.211 |
